# A novel quantile regression approach for eQTL discovery

**DOI:** 10.1101/070052

**Authors:** Xiaoyu Song, Gen Li, Iuliana Ionita-Laza, Ying Wei

## Abstract

Over the past decade, there has been a remarkable improvement in our understanding of the role of genetic variation in complex human diseases, especially via genome-wide association studies. However, the underlying molecular mechanisms are still poorly characterized, impending the development of therapeutic interventions. Identifying genetic variants that influence the expression level of a gene, i.e. expression quantitative trait loci (eQTLs), can help us understand how genetic variants influence traits at the molecular level. While most eQTL studies focus on identifying mean effects on gene expression using linear regression, evidence suggests that genetic variation can impact the entire distribution of the expression level. Indeed, several studies have already investigated higher order associations with a special focus on detecting heteroskedasticity. In this paper, we develop a Quantile Rank-score Based Test (QRBT) to identify eQTLs that are associated with the conditional quantile functions of gene expression. We have applied the proposed QRBT to the Genotype-Tissue Expression project, an international tissue bank for studying the relationship between genetic variation and gene expression in human tissues, and found that the proposed QRBT complements the existing methods, and identifies new eQTLs with heterogeneous effects genome-wideacross different quantile levels. Notably, we show that the eQTLs identified by QRBT but missed by linear regression are more likely to be tissue specific, and also associated with greater enrichment in genome-wide significant SNPs from the GWAS catalog. An R package implementing QRBT is available on our website.

## Introduction

Genome-wide association studies (GWAS) have led to remarkable progress in our understanding of the role of genetic variation in complex human diseases, resulting in the identification of thousands of common genetic variants affecting human diseases and other complex traits. Most genetic variants discovered through GWAS are non-coding, and therefore may play a role in regulating gene expression levels. Identifying genetic variants that influence the expression level of a gene, i.e. expression quantitative trait loci (eQTLs), is essential to interpreting the GWAS loci and understanding how genetic variants influence traits at the molecular level. In addition, eQTL discovery by itself is an important area, since it helps understand how genetic variants influence gene regulation and discover complex gene regulatory networks. An important resource for eQTL discovery is the Genotype-Tissue Expression (GTEx) project, a major international project designed to establish a comprehensive data resource on genetic variation, gene expression and other molecular phenotypes across multiple human tissues [1].

Most of the existing eQTL studies focus on identifying mean effects, or associations between genotype and the mean value of the expression level of a gene. However, the entire distribution of gene expression may be regulated by genetic variants. For a concrete example, variant rs7202116 at the FTO locus has been shown to be associated not only with the mean but also with the variability of body mass index (BMI) [2]. In addition, recent studies noted that heterogeneity is also associated with interactions among genetic variants (epistasis) or between variants and environment *(G × E)* [3], and hence heterogeneity can be used as a screening tool for such interactions.

For these reasons, there has been increasing attention in recent eQTL studies to quantify genetic associations at higher orders of the expression levels. Most of them focus on identifying variance eQTLs by testing heteroskedasticity, for example (1) Levene’s test [4], (2) Brown-Forsythe test [5], and (3) correlation least squared (CLS) test [6]. Both Levene and Brown-Forsythe tests test the marginal variance differences between two and more groups. While beneficial for experimental studies, their inability to account for continuous covariates such as imputed single nucleotide polymorphisms (SNPs) and principal components of population stratification largely limits their application to genetic studies in human populations. The CLS test is a regression based test. It regresses gene expression levels against genotypes, and then uses Spearman rank correlation to assess whether the residuals are heteroskedastic across genotypes. The regression based CLS method is flexible and can incorporate confounders, but the method is restricted to a family of location-scale models, where both the mean and variance of the gene expression are linear in genotypes. More recently, a Bayesian test[7] has been proposed to relax the linear assumption at the expense of increased computational cost, which could be undesirable for genome-wide identification of eQTLs that involves hundreds of millions of tests.

In addition, mean and variances alone are insufficient to describe the distributional heterogeneity. Quantile regression, proposed by Koenker and Bassett [8], has emerged as an important statistical methodology. It offers a systematic strategy for examining how covariates influence the entire response distribution by estimating various conditional quantile functions. In this paper, we extend the rank-score inference [9] in quantile regression to identify eQTLs that have impact on the gene expression distributions. The resulting quantile test, which we call Quantile Rank-score Based Test (QRBT) throughout the paper, enjoys the following advantages: (1) it is computationally efficient; (2) it can easily accommodate continuous or discrete covariates; (3) it accommodates a wide range of distributions without assuming an a priori parametric likelihood for the gene expressions; (4) it is robust to outliers in the data; (5) it simplifies the preprocessing normalization procedure; and (6) it is conservative in controlling type I errors.

We apply the proposed QRBT approach to the up-to-date Genotype-Tissue Expression (GTEx) project data [1]. The existing eQTLs identified in GTEx are based on linear regressions [10]. Our approach complements these existing studies; it leads to new eQTL discoveries that are more tissue specific, and that show higher enrichment in genome-wide significant SNPs. The results suggest that the proposed QRBT has great potential to identify disease-linked eQTLs.

## Method

### Overview of GTEx Data

We analyzed the GTEx midpoint v6 data freeze, which comprises RNA sequencing (RNA-seq) data from 7051 samples from 449 individuals representing 44 tissues (db-GaP accession number phs000424.v6.p1). We identified eQTLs separately for 4 tissues with sufficient sample sizes (*n* > 275) including: muscle-skeletal (n=361), whole blood (n=338), lung (n=278) and thyroid (n=278). Because of the relatively small sample sizes, we focused on identifying eQTLs within ± 1MB of the transcriptional start site (TSS) of each gene.

In this paper, we use genes defined as protein coding in the GENCODE version 19 [11]. The quantile normalized gene-level expression values were used for analysis as in previous studies [10] (note however that our proposed approach makes no parametric assumption for the underlying distribution of gene expression). We use the same quality control procedures as in the GTEx study [10] for consistency. We remove genes with more than 10% zero read count, as in such a case the Gaussian assumption in linear regression is violated, and also our analyses found that the existing variance eQTL method CLS [6] had largely inflated type I error. We also correct for known and inferred technical covariates including gender, genotyping array platform (Illumina’s OMNI 5m or 2.5M array), 3 principal components of SNPs and 35 PEER factors [12] of the top 10,000 expressed genes in each tissue in the analysis. More information about the preprocessing procedure of the GTEx data can be found online at http://www.gtexportal.org.

### Tissue Specific Quantile Analysis for eQTL Discovery

#### Notations and Settings

Suppose the data consist of *n* subjects who have their gene expression measured on a total of *K* genes, and are genotyped for a total of *M* SNPs. We then denote **Y** as a *n × K* gene expression matrix, where *Y_ik_* is the gene expression level of the *i*-th subject on the *k*-th gene, *G_k_*. We denote X as a *n × M* genotype matrix, where *x_i,j_* is the *i*-th subject’s genotype on the *j*-th SNP. We finally denote z_*i*_ as the vector of covariates of the *i*th subject, including the intercept. Throughout the paper, we denote 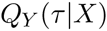 as the τ-th conditional quantile of *Y* given *X*.

Let Λ_k_ be the subset of SNPs that are within ±1MB of the TSS of gene *G_k_*, then for each SNP-gene pair *(j, k)* where 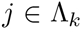 and 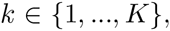 we build the following linear quantile model

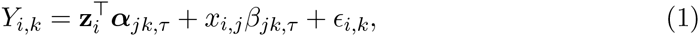

where ϵ*_i,k_* is the random error whose τ-th conditional quantile 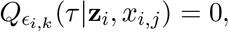 and τ ∈ (0,1) is the quantile level of interest. Under Model (1), the conditional quantile of *Y_i,k_* is a linear function of z_i_ and 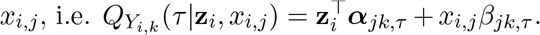. In this model, β_*jk*,τ_ is the primary parameter of interest, which characterizes the association between the genotype *x_i,j_* and the gene expression level of *G_k_*. The goal of the analysis is to identify the *(j, k)* pairs whose 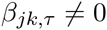 for any given t τ ϵ (0,1).

#### Quantile Rank-score Based Test at a Fixed Quantile

At a fixed quantile level, the existing inference tools for quantile regression can be generally classified into three categories: Wald-type inference, rank-score method and resampling methods [13]. The Wald-type inference requires the direct estimation of the asymptotic variance-covariance matrix. That, however, is computationally difficult, since the limiting variance-covariance matrix contains the density of the error ϵ_i,k_ at the τ-th quantile. In the framework of quantile regression, the error distribution is non-i.i.d. and completely unspecified. As a result, the limiting variance-covariance matrix contains *n* nuisance parameters. Without a parametric likelihood, it is hard to estimate those local densities. Several kernel based approaches have been proposed in this context, but their estimates are often unreliable at extreme quantiles or with relatively small sample sizes. In our preliminary analyses, we also found that direct Wald type inference with kernel estimated densities has inflated type I errors at very small significance level (e.g. *α* < 1e — 6). Alternatively, resampling based inference such as bootstrap does not require density estimation; however it is computationally intensive, and hence undesirable in GTEx applications where one needs to repeat the analysis for hundreds of millions of SNP-gene pairs for each tissue.

We hence propose to extend the rank-score test [9] for eQTL discovery. For any fixed quantile τ, the rank score function in quantile regression can be written as

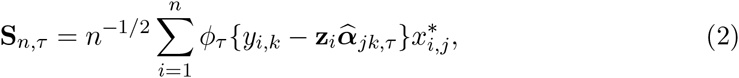

where 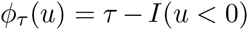 is an asymmetric sign function, and 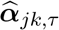 is the estimated coefficient under the null *H_0_*: β_jk,τ_ = 0. Define 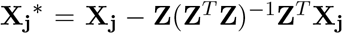 as the residual vector of **X_j_** projected on the column space of **Z** (the design matrix under the null), then 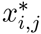 in (2) is the ith element of 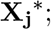 the projection is done to achieve the asymptotic independence between **X** and **Z**. Hence the test statistics S_n,τ_ measures the quantile association between **Y**and **X** that is accounted for the co-linearity between **X** and **Z**. Since the function 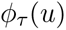 essentially measures the signs of the residuals, S_n,τ_ is in the category of rank-based statistics, and hence also called rank score function.

Note that 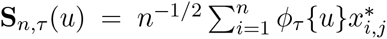 is the quantile regression estimating functions that is associated with β_*jk*,τ_. When *u* is the residual under the null hypothesis, **S**_n,τ(*u*)_ is close to zero if and only if the null hypothesis is true. Any deviation from the null model will push S_n_,_τ_(u) away from zero. Consequently, one could construct atest statistics to test whether β_*jk*,τ_ = 0 by

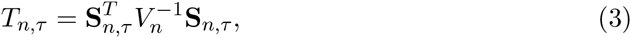

where 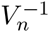 is the variance of **S***_n,τ_* such that 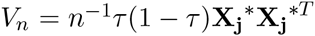. According to the rank-score inference [9],

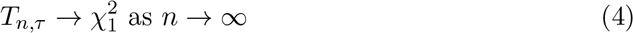

under the null hypothesis β_*jk*,τ_ = 0. Similar construction in maximum likelihood estimation (MLE) is called generalized likelihood ratio statistics [14].

The asymptotic distribution of Equation (3) was established under the assumption of i.i.d. errors. Although this assumption is often unrealistic for quantile regressions, many studies [15, 16] have consistently found that the rank score test is very robust with non-i.i.d. errors. A generalized rank score test with non-i.i.d. densities could be found in [16]. However, it again requires the estimation of the nuisance parameters 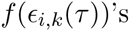. Even though it is theoretically appealing, the generalized rank score test is much harder to implement. For this reason, we will investigate the performance of the simple rank score test (2) in the setting of eQTL discovery. The quantile regression rank-score test enjoys the following advantages. (1) It is a distribution-free statistic. Under the framework of quantile regression, the test does not assume any likehood distributions on the gene expressions. Hence it can be applied to any gene expression data without requiring a pre-transformation to achieve normality. (2) The construction of the test statistics is simple and avoids the estimation of local densities. Although the asymptotic theory assumes an independent and identically distributed (i.i.d.) error model, the rank score test has very robust performance under various error structures and distributions. (3) It is computationally fast. To construct rank-score test statistics, we only need to estimate the null model where β_*jk*,τ_ = 0, which greatly reduces the computation cost from *M × K* pair-wise regressions for each SNP-gene pair to *K* regressions.

#### Composite Rank-score Test

Instead of individual quantile level p-values, it would be desirable to have a single pvalue for a SNP-gene pair from a composite test across multiple quantile levels. Suppose we consider ℓ quantile levels of *t_1_,t_2_, &,tℓ,* then define 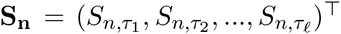 as the vector of rank score test statistics at the corresponding quantile levels. We can show that, under the null hypothesis, **S_n_** asymptotically follows a multivariate normal distribution,

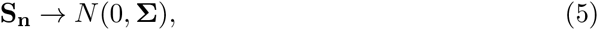

where Σ is the ℓ × ℓ variance-covariance matrix. The diagonal elements of Σ are 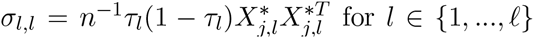, and the off-diagonal elements of Σ are 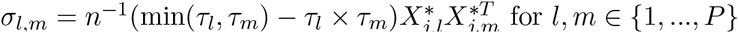 and *l ≠ m*.

A natural composite rank score test statistic can be constructed by the following quadratic form in **S_n_**:

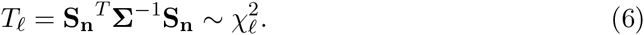

To select the quantile levels, one could either choose *l* evenly spaced quantile levels, or go with the commonly used quantile levels, such as 0.1, 0.25, 0.5, 0.75 and 0.9. Depending on the nature of the application, one may also select quantile levels in a specific interval of interest. For example, if we are only interested in identifying eQTLs that are associated with extreme values of gene expression, we could select only quantiles at the upper tail.

The composite rank score *T_ℓ_* combines the quantile associations over multiple quantiles, regardless of the directions of the quantile associations. To some extent, one can view the mean effect as 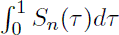, an integrated quantile effect. When the quantile association is homogeneous at all the quantiles in terms of both direction and magnitude, then testing the composite quantile association at £ evenly spaced quantile levels is equivalent to testing the mean effect. When the association is heterogeneous across quantile levels, especially when the association is “crossing” over quantile levels, i.e. **S_n_** is positive for certain quantiles but negative for others, or the association only manifests at extreme quantiles, the linear regression could underestimate, or even completely miss the underlying SNP-gene link. The composite quantile test hence has better chance to discover such heterogeneous associations. As we report below in the Results section, the eQTLs associated with heterogeneous associations are more likely to be associated with complex traits, which underscores the potential of quantile analysis in eQTL discovery.

## Results

### Comparison methods

Here we present a simulation study to validate the type I error of the proposed quantile test, and its application to the GTEx data to illustrate the potential value of the quantile based test. When implementing the proposed QRBT test, we considered 5 quantile levels at τ = (0.15, 0.25, 0.5, 0.75, 0.85), and combine their rank score functions to test whether genetic variants have effect on the entire distribution of gene expression levels. In both studies, we compare the proposed quantile approach to the following two existing methods: (1) linear regression (LR) following the GTEx analysis protocol, and(2) CLS test. Linear regression is the most commonly used method for eQTL discovery. It assumes that the gene expression level *y_í,k_* (after quantile-normalization [10]) follows a linear model

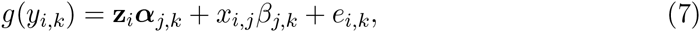

where *g*() is the quantile-normalization function, and e_i,k_ is the random error with mean zero. Here β_j,k_ measures the effect of the variant *x_i,j_* on the mean of the normalized *y_i,k_* (see the above Section on Overview of GTEx Data).

The CLS test [6] takes the residuals from the linear regression (7), and then calculates the Spearman correlation between the genotype *x_i,j_* and the residuals squares 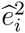. If the resulting correlation is significant, that suggests that SNP *j* is associated with the variance of the gene expression level.

### Simulations

We first investigate the type I errors of the three approaches using the GTEx gene expression data in lung tissue. Specifically, we randomly select a gene *G_k_* from all the genes in the GTEx lung tissue with non-zero expression in at least 90% of the subjects, and then randomly select a SNP *j* from all the genotyped and imputed SNPs. We randomly permute the genotypes *x_i,j’_*s to remove any association between SNP *j* genotype and gene *k* expression level. By only permuting *x_i,j’_*s, we preserve the association between phenotype and covariates. We then apply all the three approaches to test the conditional association between *y_í,k_* and permuted *x_i,j_*.

The type I errors estimated from 1 billion Monte-Carlo replicates are presented in Table 1 at multiple significance levels ranging from 0.05 to as low as 10^-7^. As shown all the approaches under consideration have well-controlled type I errors, with the proposed QRBT being slightly more conservative.

**Table 1:**
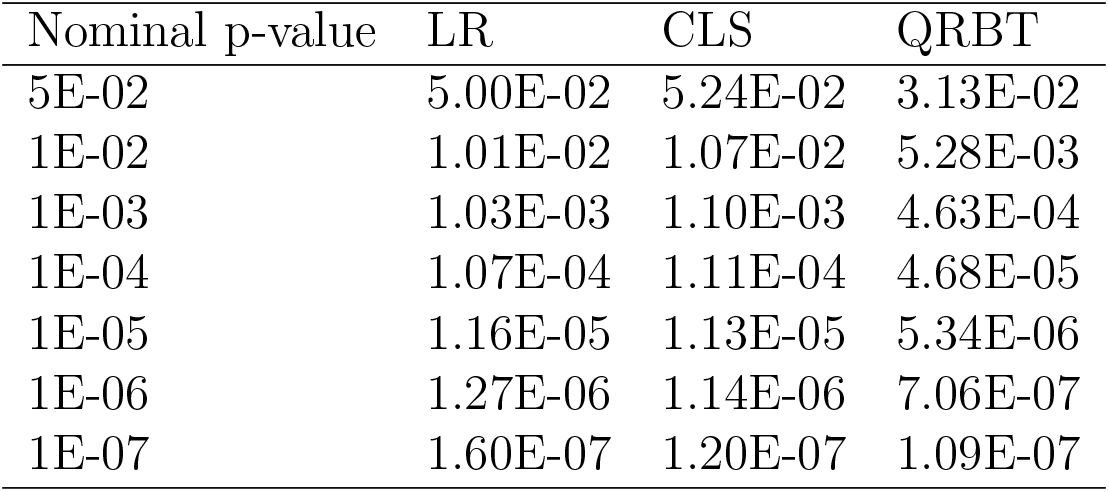
Type I error of three approaches in simulations based on the lung tissue data.

### GTEx Data Analysis

#### eQTLs identified in four tissues

Supplemental Table S1 provides information for each of the four tissues we analyzed (muscle-skeletal, whole blood, lung and thyroid), including the sample size, the number of genes with < 10% zeros, the number of SNPs genotyped or imputed within the ±1MB neighborhood of the genes, the number of SNP-gene pairs and the p-value threshold needed to control the family-wise error rate (FWER) at the 5% level with Bonferroni correction.

Figure 1 presents the Venn diagrams of identified SNP-gene pairs using LR, CLS and QRBT in four tissues controlling for 5% FWER. The patterns in all four tissues are similar. In particular, LR identified the most significant eQTLs, CLS identified the least, and QRBT in between. This suggests that linear regression remains a powerful tool to identify eQTLs, while the CLS test may have limited power in eQTL applications. The eQTLs identified by QRBT overlap to a large extent with those identified by LR; however there is a large number of eQTLs uniquely identified by QRBT. A careful examination on quantile specific effects reveals that most of the overlapping eQTLs have homogenous effects across the quantile levels. In fact LR is expected to be more powerful than QRBT under the assumption of homogeneous association due to its parametric assumption. In contrast, the eQTLs that are uniquely identified by QRBT often exhibit substantial heterogeneity across the quantiles, and consequently are missed by linear regression. To illustrate the differences between the two sets of SNPs (uniquely identified by QRBT vs. those identified by both LR and QRBT), we quantify the degree of heterogeneity for each SNP-gene pair as the log transformed ratio between the standard deviation and the mean of their 5 estimated quantile coefficients β_jk_,_τ_s. In Figure 2, we plotted overlayed histograms of the resulting heterogeneity indexes between the two sets of SNPs. As shown, the eQTLs that are uniquely identified by QRBT presented more heterogeneous effects compared to those identified by both LR and QRBT.

**Figure 1:**
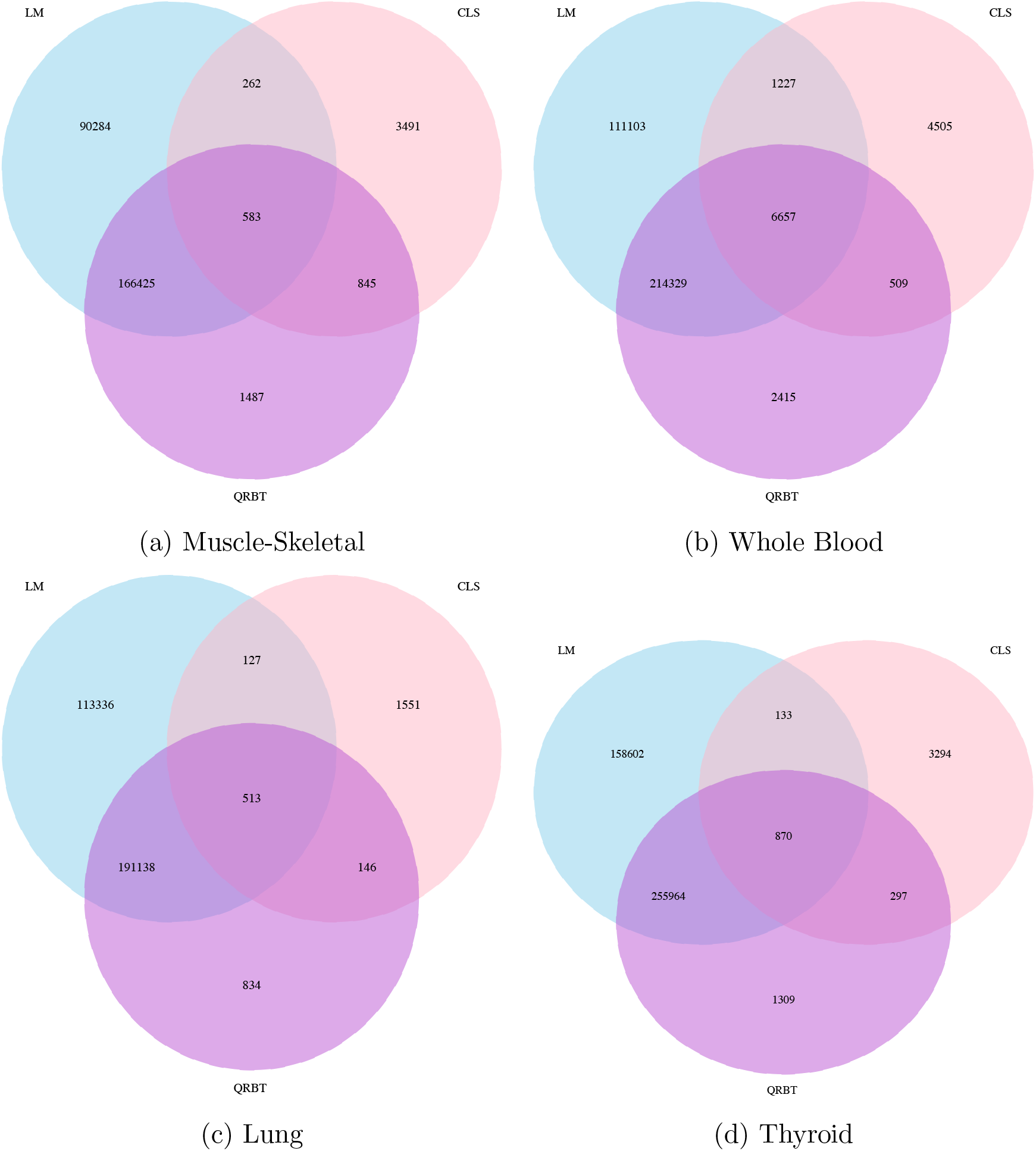
Venn diagram depicting overlap among SNP-gene pairs identified by LR, CLS and QRBT controlling FWER at *α* = 0:05.

**Figure 2:**
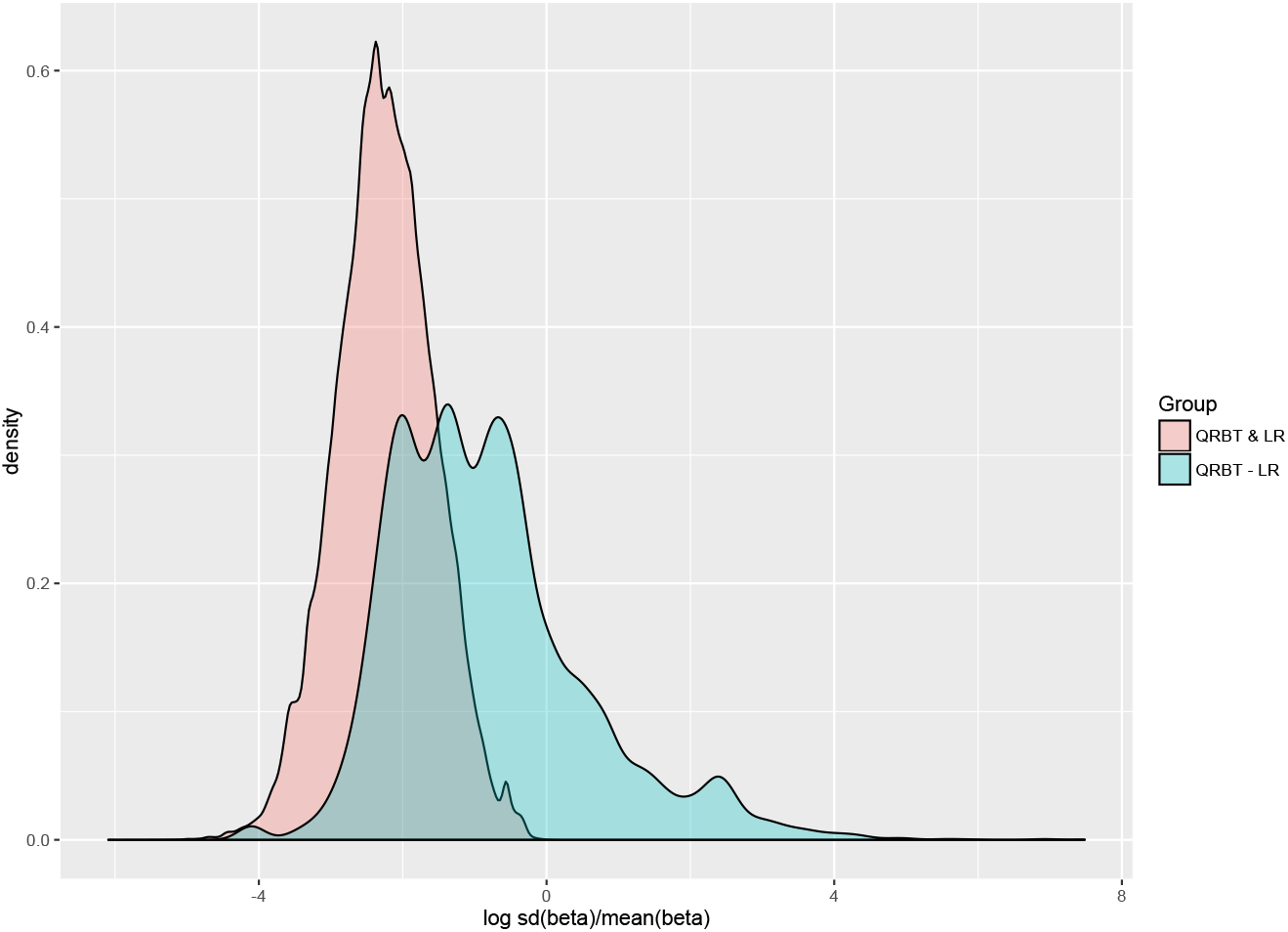
Histogram of the log transformed ratio between the standard deviation and the mean of their 5 estimated quantile coefficients β_*jk*,τ_s in four tissues. The pairs identified by QRBT but missed by LR tend to be more heterogeneous than the pairs identified by both methods.

**Figure 3:**
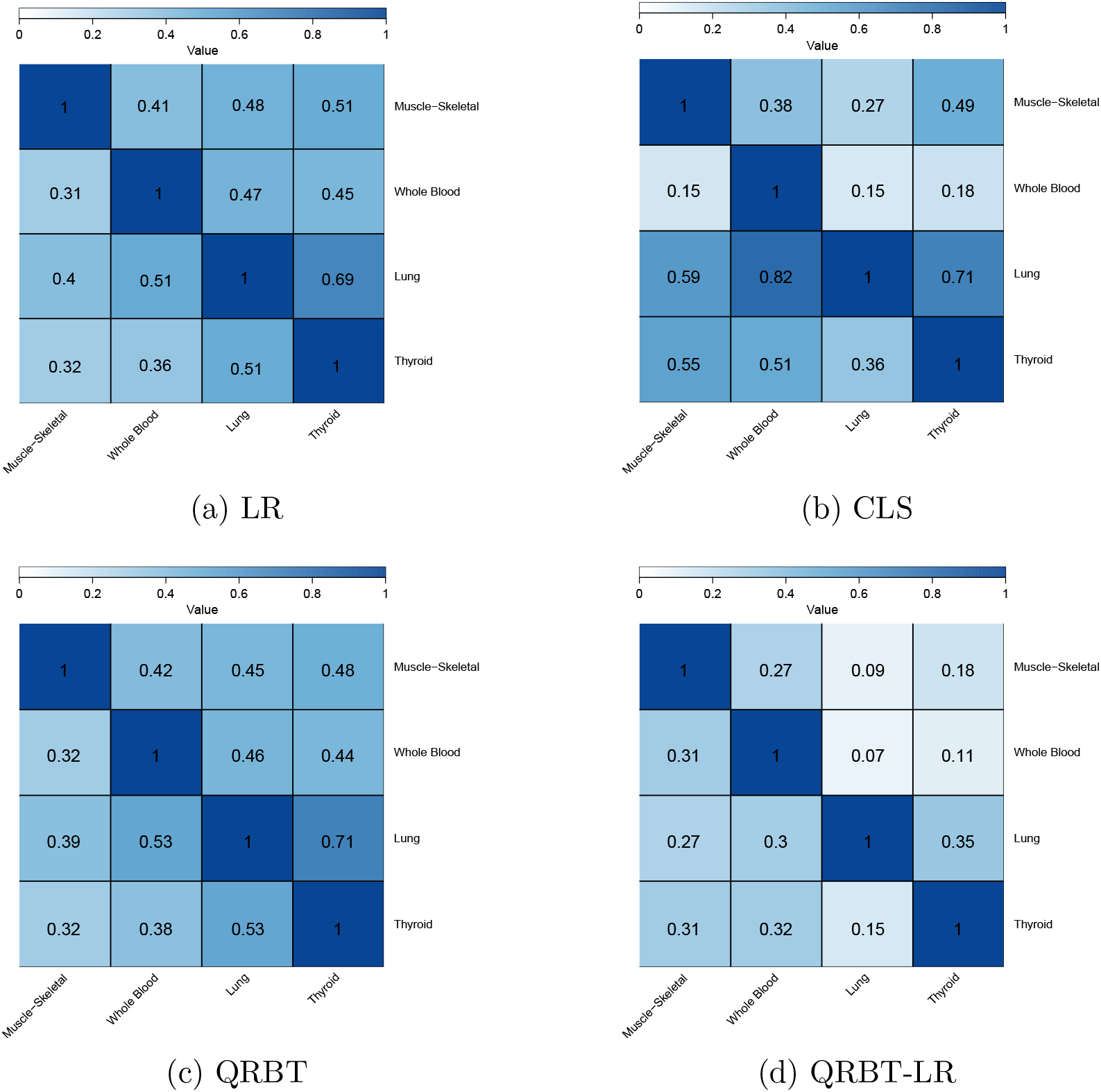
Cross-tissue sharing of eQTLs. The entry in row *i* and column *j* is an estimate of π*_ij_* = Pr(eQTL in tissue *i* | eQTL in tissue *j*). QRBT-LR has the lowest levels of eQTL sharing with other tissues.

#### Explore the eQTL Association Patterns using quantile specific QRBT

To get a better understanding of the differences in the eQTLs identified by the different methods, we looked at the association patterns of those identified eQTLs. One advantage of quantile based approach is to investigate how the eQTLs impact the entire distribution of the gene expression. To do that, we estimate the quantile coefficients on a fine grid of quantile levels (49 evenly spaced quantile levels ranging from 0.02 to 0.98). In Figure 4, we plotted the estimated conditional distribution functions of gene expression levels with different genotypes in selected pairs in thyroid tissue. Specifically, the black curve is the estimated quantile function with reference SNP values, while the red and green curves are the estimated quantile functions with one or two alternative alleles assuming additive genetic models.

**Figure 4:**
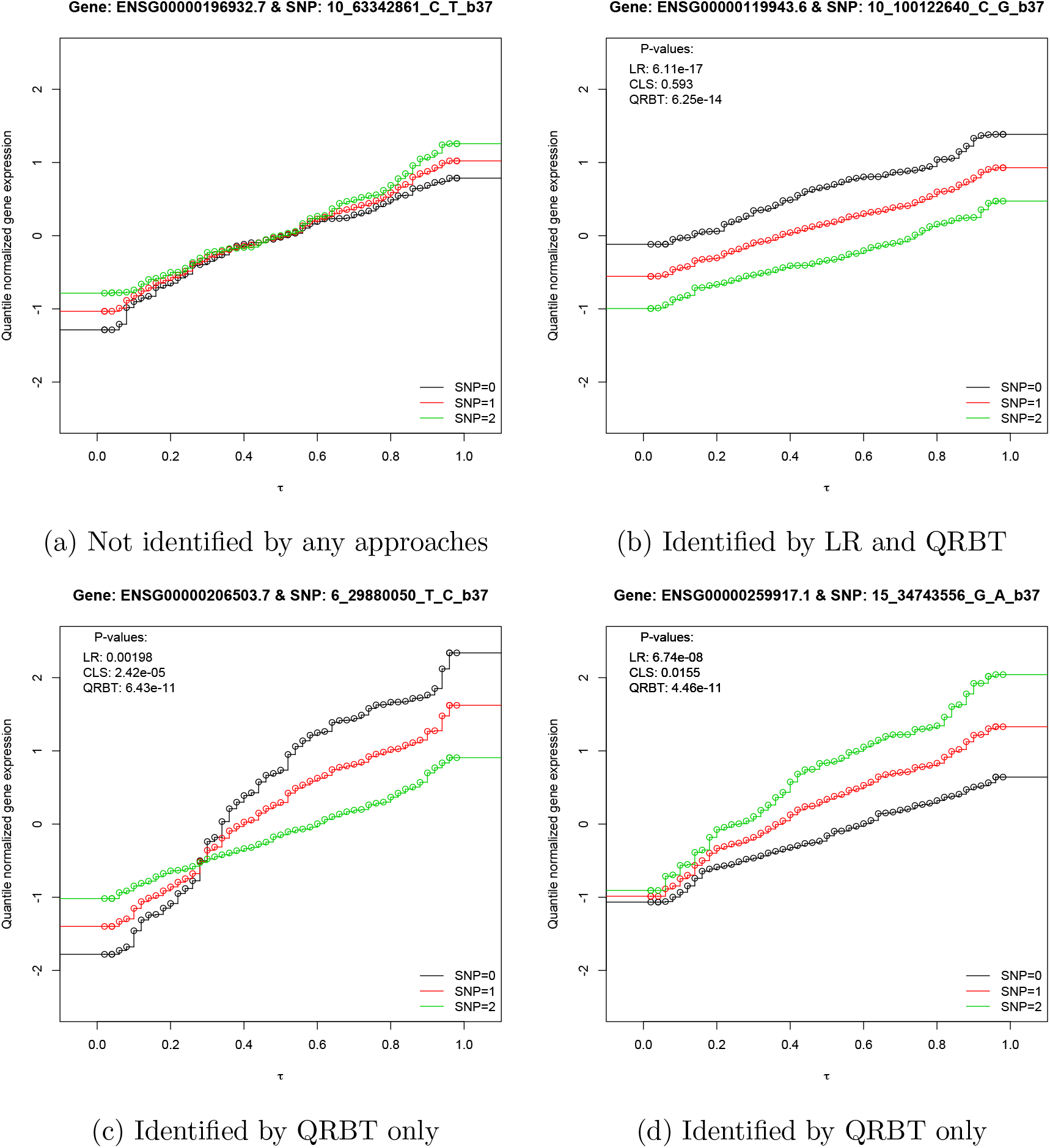
The estimated conditional distribution functions of gene expression levels for a few SNP-gene pairs in thyroid tissue. The x-axis is the grid of quantile levelsr τ ϵ (0,1) and the y-axis is the estimated conditional distribution functions for each quantile level given three SNP values and averaged covariates. This figure presents how the entire distribution of gene expression differs by SNP values for 4 SNP-gene pairs.

Each sub-figure represents a distinctive association pattern. Figure 4a presents a SNP-gene pair that is not identified by any of the approaches. As shown, the three curves are nearly identical at all the quantile levels, which suggest that the SNP genotype has little impact on the gene expression level. As expected, none of the approaches identify it as an eQTL. Figure 4b presents a SNP-gene pair that is identified by both LR and QRBT, but missed by CLS. In this case, the effect of the SNP on gene expression is homogeneous in both the direction and magnitude across all quantile levels. In this case, LR is more efficient than QRBT with smaller p-values. Figure 4c presents a SNP-gene pair with a “crossing” heterogeneous effect such that the SNP promotes the gene expression at lower quantiles, but suppress the gene expression at upper quantiles. Such eQTLs would be missed by LR as their effect at lower and upper quantiles cancels out at the mean level; in contrast, the proposed QRBT is not affected by such crossing effect because the test statistics accumulates the squared estimating functions. As shown in their p-values, the CLS test detects such association pattern with a much limited power. Finally, Figure 4d presents another heterogeneous effect pattern, in which case the SNP has an effect that is mostly evident at upper quantile levels. In this case the SNP has an effect only at upper quantile levels, and LR misses the local effect while QRBT captures it.

These examples illustrate the advantage that QRBT can have over LR in identifying SNP-gene pairs with heterogeneous effects, and in providing a more comprehensive association picture for eQTL discoveries.

#### Tissue-specific effects in the four tissues

We have also investigated the sharing patterns of eQTLs across tissues, for each method separately. As complex traits are assumed to be influenced by regulatory elements that act in a tissue-specific manner, tissue-specific eQTLs are more likely to be linked with disease risk than cross-tissue eQTLs [17]. To understand the eQTLs sharing patterns for each method, we compute a pairwise eQTL sharing estimate π*_ij_* = Pr(eQTL in tissue *i* | eQTL in tissue *j*). In Figure 3 we show the pairwise eQTL sharing π*_ij_* for the different approaches. We denote by QRBT-LR the eQTLs from the SNP-gene pairs identified by QRBT but missed by LR in the same tissue. In multi-tissue results, QRBT-LR are the eQTLs from the SNP-gene pairs identified by QRBT but missed by LR in at least one tissue. As shown, eQTLs that are uniquely identified QRBT are the least shared in all approaches considered.

In Table 2 we show the relative risk (RR) of being tissue-specific eQTLs for eQTLs identified by each approach in comparison with LR. Out of the 6705 eQTLs that were identified by QRBT but not linear regression, 89% of the eQTLs that tissue specific. In comparison, only 54% LR-identified eQTLs are tissue specific. Statistical test on the relative risks also show that CLS, QRBT, QRBT-LR are all significantly more likely to detect tissue-specific eQTLs than LR, the eQTLs identified by QRBT-LR, however, are most likely to be tissue-specific (RR: 1.65; 95% CI (1.63, 1.65)).

**Table 2:** The tissue-specificity of eQTLs identified by different approaches. The relative risk (RR) is calculated as the probability of being tissue-specific eQTLs for eQTLs identified by each approach in comparison with LR. The eQTLs identified by QRBT-LR are most likely to be tissue-specific.

|  | LR | QRBT | CLS | QRBT-LR |
| --- | --- | --- | --- | --- |
| No. of eQTLs | 470413 | 310931 | 15828 | 6705 |
| % of tissue-specific eQTLs | 54% | 57% | 75% | 89% |
| RR | ref | 1.05 | 1.37 | 1.64 |
| 95% CI | ref | (1.04, 1.05) | (1.36, 1.39) | (1.63, 1.65) |
| P-value | ref | 5.8E-112 | < 2.2E-308 | < 2.2E-308 |

#### Enrichment of GWAS SNPs among the eQTLs identified in the four tissues

We investigate here the enrichment of eQTLs identified by the different methods in genome-wide significant SNPs from the GWAS catalog ([18]; version June 2016). Table 3 presents the enrichment results. The GWAS enrichment is calculated with reference to LR by the relative risk (RR), the ratio of the probability of an eQTL identified by one approach to be in the GWAS catalog relative to an eQTL identified by LR. Table 3 shows that both CLS and QRBT-LR are significantly enriched in GWAS catalog SNPs in comparison with LR, with QRBT-LR having the biggest estimate (RR: 1.74; 95% CI (1.30, 2.32)) of enrichment. Results for each separate tissue are available in Supplement Table S2.

**Table 3:** The enrichment of identified eQTLs in SNPs from the GWAS catalog in four tissues. The relative risk (RR) is calculated as the probability of being in GWAS catalog for eQTLs identified in each approach in comparison with LR. The eQTLs identified by QRBT-LR are most likely to be enriched in SNPs from the GWAS catalog. Figure 1: Venn diagram depicting overlap among SNP-gene pairs identified by LR, CLS and QRBT controlling FWER at *a* = 0.05.

|  | LR | QRBT | CLS | QRBT-LR |
| --- | --- | --- | --- | --- |
| No. of identified eQTLs | 470413 | 310931 | 15828 | 6705 |
| No. of identified eQTLs in GWAS | 1896 | 1315 | 107 | 47 |
| RR | ref | 1.05 | 1.68 | 1.74 |
| 95% CI | ref | (0.98, 1.13) | (1.38, 2.04) | (1.30, 2.32) |
| P-value | ref | 1.79E-01 | 1.34E-07 | 1.43E-04 |

## Discussion

In this paper, we develop a new quantile regression based test procedure for the genome-wide identification of eQTLs. Unlike linear models which focus on the effect of SNPs on mean expression levels, quantile regressions characterize a comprehensive picture of how genetic variants affect gene expressions at different quantiles. Test statistics are derived from the rank score function in quantile regressions. In particular, for the fixed quantile test, the test statistic is a quadratic form of the rank score at a fixed quantile. For the composite quantile test, we combine rank scores across a set of quantiles. The test statistics have explicit asymptotic distributions under the null, and thus the hypothesis testings are computationally efficient. The proposed method can easily accommodate continuous or discrete covariates, and is robust against non-i.i.d. error terms. In the simulation study, we show that the method strictly controls the type-I error. In the GTEx v6 data analysis, the proposed method not only identifies eQTLs with significant mean effect differences, but also makes many unique discoveries not obtainable from linear models. We further investigate the additional discoveries and obtain interesting patterns of how genetic variants regulate gene expressions with heterogeneity in effect across different quantiles. The GWAS enrichment analysis shows that the additional eQTLs are highly enriched in the SNPs in the GWAS catalog. Therefore those eQTLs detected by QRBT but missed by LR might be interesting in understanding the existing GWAS findings. Overall, the proposed method provides an alternative approach for eQTL detection, and the results complement the existing knowledge by understanding the differential expression across the entire distribution.

There are several interesting directions for future work. One is to better accommodate zero inflation in gene expression data. So far, we have focused on genes with fewer than 10% zero read count. In practice, many genes have excessive zero read counts due to various experimental and biological reasons. The abundance of zeros may be problematic with the lower quantiles and leads to numerical instability of the proposed method. New methods are needed to deal with the zero inflation problem. For example, one may add small perturbations to the zero values to break the ties. Conceptually this will not affect the estimation very much but will greatly improve the computational performance of the method. Another idea is to introduce an additional latent variable to indicate the presence of zeros [19], and model zeros separately. A second direction is to build joint models for eQTL analysis in multiple tissues simultaneously. It is well known that most eQTLs are shared across tissues, while some are highly tissue specific [10]. Analyzing gene expression data from multiple tissues simultaneously will increase the power of eQTL detection by borrowing strength across tissues, and also will facilitate the assessment of tissue specificity [20, 21]. However, how to extend the quantile regression method to multiple tissues is not trivial. A SNP may regulate the expression level of a gene at different quantiles in different tissues. Furthermore, the computational burden will be more severe in multi-tissue analysis. This calls for further investigation. A third direction is to use functional effect predictions for genetic variants, non-tissue specific such as GERP [22] and Eigen [23], or tissue-specific [24] as priors to improve power to identify eQTLs, especially in trans-eQTL mapping studies.
Software implementing the proposed QRBT is available as an R package on our website at https://XiaoyuSong.shinyapps.io/QRBT. The database containing eQTLs with p-value < 10^-6^ in at least one of the three approaches (LR, CLS and QRBT) as well as their summary statistics is also available on this website.

## Acknowledgments

This research is supported by NIH grant R01HG008980 and R03HG007443. IIL has been supported by NIMH grant MH106910. The GTEx Project is supported by the Common Fund of the Office of the National Institutes of Health with additional funds provided by the NCI, NHGRI, NHLBI, NIDA, NIMH, and NINDS. We would also like to thank two students Xianling Wang and Zhengwei Zhou for help with data presentation.

## Web Resources

GTEx: http://commonfund.nih.gov/GTEx, and http://www.gtexportal.org GWAS catalog: https://www.ebi.ac.uk/gwas GENCODE version 19: http://www.gencodegenes.org/releases/19.html

## Supplementary Figure

**Table S1:**
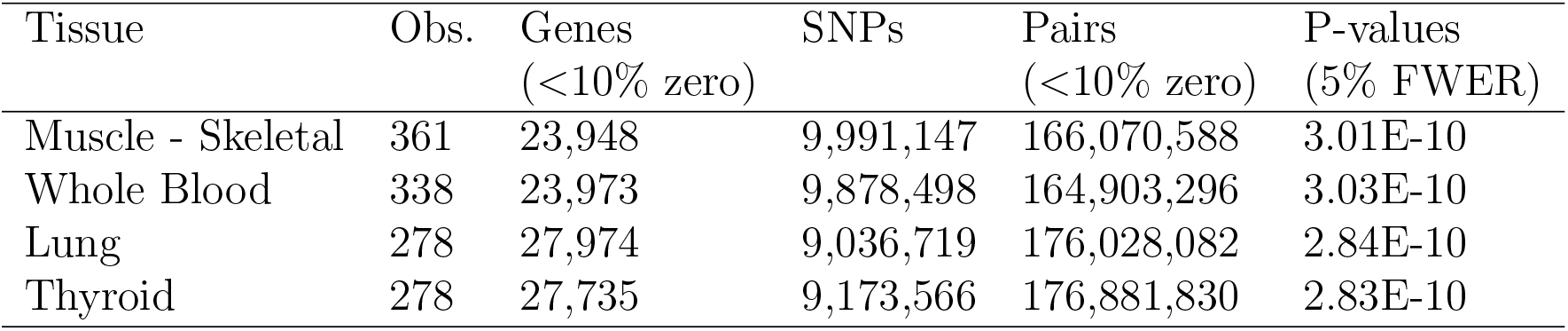
Description of the GTEx data in four tissues.Table S2: The enrichment of identified eQTLs in GWAS catalog in each tissue. The relative risk (RR) is calculated as the probability of being in GWAS catalog for eQTLs identified in each approach in comparison with LR.

**Table S2:** The enrichment of identified eQTLs in GWAS catalog in each tissue. The relative risk (RR) is calculated as the probability of being in GWAS catalog for eQTLs identified in each approach in comparison with LR.

|  | LR | QRBT | CLS | QRBT-LR |
| --- | --- | --- | --- | --- |
| **Muscle-skeletal** |  |  |  |  |
| No. of identified eQTLs | 176888 | 118898 | 5000 | 2213 |
| No. of identified eQTLs in GWAS | 764 | 517 | 39 | 22 |
| RR | ref | 1.01 | 1.81 | 2.30 |
| 95% CI high | ref | (0.90, 1.13) | (1.31, 2.49) | (1.51, 3.51) |
| P-value | ref | 9.06E-01 | 2.51E-04 | 7.00E-05 |
| **Whole Blood** |  |  |  |  |
| No. of identified eQTLs | 208620 | 137404 | 11568 | 2868 |
| No. of identified eQTLs in GWAS | 957 | 690 | 92 | 17 |
| RR | ref | 1.09 | 1.73 | 1.29 |
| 95% CI | ref | ( 0.99, 1.21) | (1.40, 2.15) | (0.80, 2.08) |
| P-value | ref | 6.93E-02 | 3.10E-07 | 2.92E-01 |
| **Lung** |  |  |  |  |
| No. of identified eQTLs | 202451 | 126707 | 2302 | 977 |
| No. of identified eQTLs in GWAS | 861 | 525 | 23 | 4 |
| RR | ref | 0.97 | 2.35 | 0.96 |
| 95% CI | ref | (0.87, 1.09) | (1.56, 3.55) | (0.36, 2.57) |
| P-value | ref | 6.37E-01 | 2.97E-05 | 9.39E-01 |
| **Thyroid** |  |  |  |  |
| No. of identified eQTLs | 275495 | 174387 | 4467 | 1606 |
| No. of identified eQTLs in GWAS | 1141 | 718 | 40 | 10 |
| RR | ref | 0.99 | 2.16 | 1.50 |
| 95% CI | ref | (0.91, 1.09) | (1.58, 2.96) | ( 0.81, 2.80) |
| P-value | ref | 9.01E-01 | 8.51E-07 | 1.95E-01 |

